# The impact of the serotonergic psychedelic DOI on active vision in freely moving mice

**DOI:** 10.1101/2025.10.14.682230

**Authors:** Rolf J. Skyberg, Christopher W. Fields, Dylan M. Martins, Cristopher M. Niell

## Abstract

Psychedelics offer a unique opportunity to investigate the neural mechanisms governing both normal and altered perceptual states, yet relatively little is known about their impact on visual processing at the level of neural coding, particularly in the context of active vision. Here we examined how the serotonergic psychedelic DOI (2,5-dimethoxy-4-iodoamphetamine) influences neural activity within the visual cortex (V1) of freely moving mice engaged in naturalistic vision. Previous work has demonstrated that, under normal conditions, gaze shifts trigger a temporal sequence in V1 that encodes visual information in a coarse-to-fine sequence. Here we found that DOI had a diverse and temporally dynamic impact on response amplitudes across this sequence — while net population-level firing was modestly suppressed, individual neurons showed large increases or decreases in their gaze shift responses. Strikingly, this bidirectional modulation was correlated with gaze-shift response latency. DOI had mixed effects on short latency, low spatial frequency preferring neurons, but became increasingly suppressive across the sequence, ultimately suppressing nearly all long latency, high spatial frequency preferring neurons. These findings demonstrate how psychedelics disrupt naturalistic visual processing and support a framework in which visual hallucinations result from an imbalance in coarse and fine input integration during active vision.

**Highlights:**

- We recorded neural activity from V1 of freely moving mice before and after administration of the psychedelic DOI.
- DOI had diverse effects on visual response amplitude following a gaze shift, with some neurons increasing while others decreased.
- The modulation of responses was temporally dynamic, shifting across the response sequence.
- Longer latency neurons, which preferentially encode high spatial frequency information, were more consistently suppressed
- Together these effects result in a disruption of population response and coarse-to-fine processing, providing potential mechanisms for altered visual perception

## Introduction

Psychedelics, a subclass of hallucinogenic compounds, have the ability to disrupt perception, cognition and mood^1^, often resulting in profound and highly meaningful experiences. Recently, there has been a resurgence of interest and use of psychedelics for their potential therapeutic application for conditions including treatment-resistant depression, anxiety and trauma-related disorders^2^. Beyond their long-term therapeutic and neuroplastic effects^3,4^, psychedelics also induce acute perceptual effects, altering sensory processing in a way that generates perceptual distortions and hallucinations^1,5^. Determining the neural mechanisms underpinning these sensory disruptions remains an area of active exploration^6^ with potential to deepen our understanding of how the brain creates perception in normal and psychedelic-induced states.

The psychedelic compound DOI (2,5-dimethoxy-4-iodoamphetamine) is a serotonergic agonist that is known to be a powerful hallucinogen in humans^1^. In animal models, DOI induces a variety of serotonergic-dependent behaviors such as the head-twitch response^7^, dose-dependent changes in locomotor activity^8^ and forepaw stereotypy^9^. At the neurophysiological level, DOI has been shown to alter sensory processing in the auditory cortex^10^ and visual cortex (V1) of awake head-fixed mice^11^, attenuating the bottom-up stimulus-evoked responses in both primary sensory cortices. Recent work using functional magnetic resonance imaging has also revealed psychedelic-induced suppression of visually evoked activity in V1 of human participants engaged in a visual task^12^. Thus, attenuated stimulus-evoked activity is a central feature of psychedelic’s effect on sensory processing.

More recently, there have been efforts to study visual processing in more ethologically relevant contexts such as free movement^13–17^. Importantly, these studies allow for the natural visual behaviors of the animal such as eye and head movements, which are prevented during head-fixed paradigms. Notably, we recently demonstrated a temporal sequence of coarse-to-fine processing that is initiated by saccade-like gaze-shifting head and eye movements in the mouse visual cortex^16^, highlighting the importance of studying sensory function in naturalistic contexts such as free movement. It is unclear how the effects of DOI discovered using head-fixed paradigms translate to naturalistic contexts that allow for active sampling behaviors, and in particular, how it impacts gaze shift evoked responses dynamics. To address this we studied the impact of DOI on both active visual sampling behaviors and neural activity in V1 of awake, freely moving mice. This provided a richer view of the impact on visual responses, revealing that it was both diverse and temporally dynamic, thereby shifting neural coding and the balance of coarse-to-fine processing.

## Results

To investigate the effects of DOI on natural visual processing in freely moving mice, we used a head-mounted recording system consisting of a camera to record the position of the right eye, a forward-facing wide-angle (120°) camera to capture the visual scene, an inertial measurement unit (IMU) to record head angular velocity, and a chronically implanted 128-channel linear silicon probe in V1 for single-unit electrophysiology (Figure 1A). Mice were first placed into the freely-moving arena containing black and white blocks and for ∼45 minutes before being head-fixed on a spherical treadmill for ∼30 minutes to measure visual tuning properties using traditional computer-generated stimuli. Mice were then given a subcutaneous injection of either DOI (10mg/kg in saline) or saline (0.9% NaCl) and the experiments were repeated in reverse order (Figure 1B). A camera above the recording arena allowed us to track the position of the animal during the free movement portions of the recording (Figure S1A).

**Figure 1.**
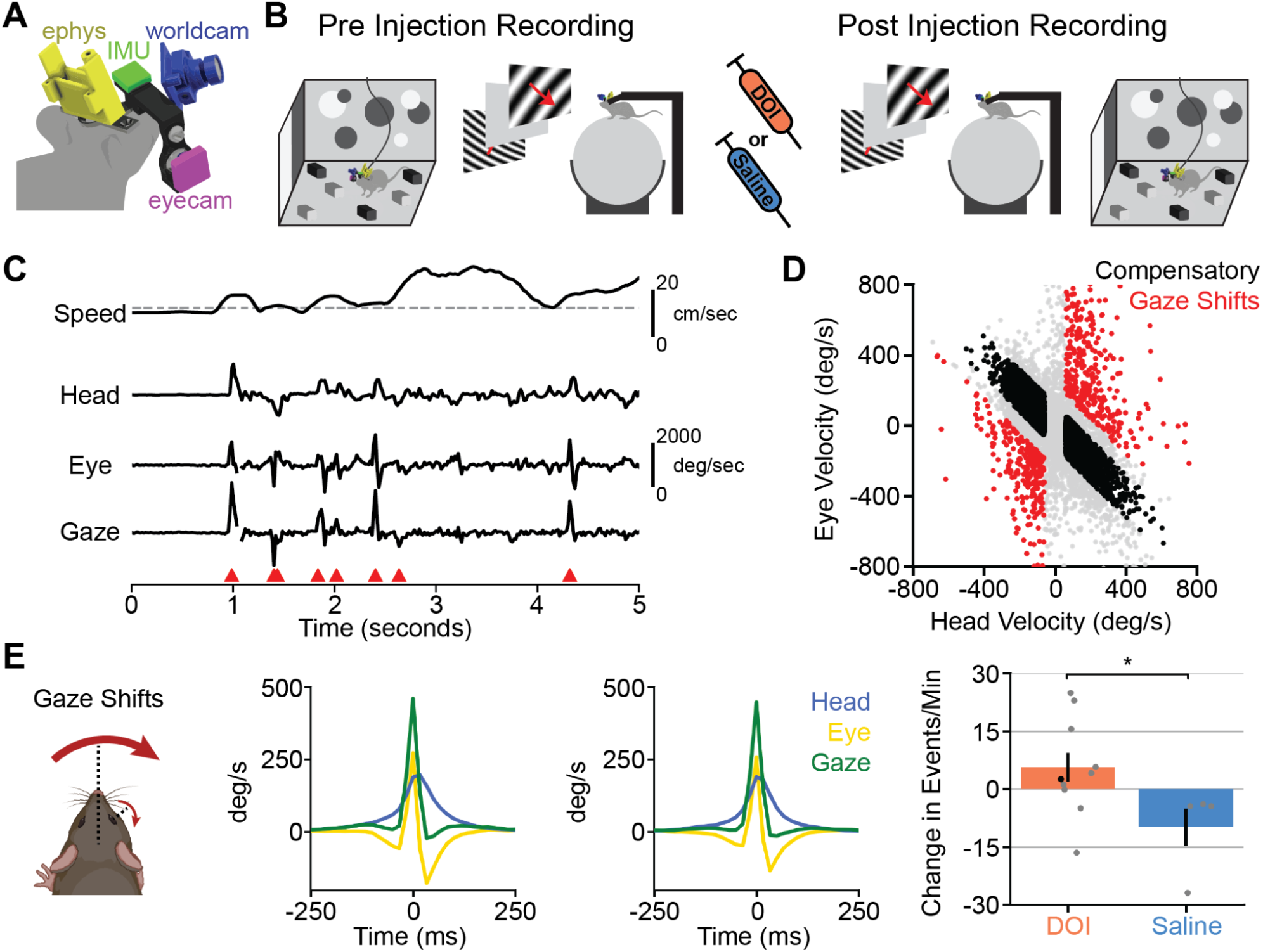
Experimental setup and effect of DOI on active sampling behaviors. (A) Schematic of head-mounted recording system including 128-channel silicon electrode implanted in V1, eye camera, world camera and inertial measurement unit (IMU). (B) Experimental design to record neural responses during free movement and head-fixation before and after administration of either DOI (n = 10) or saline (n = 4). Stimuli shown during head-fixation included reverse checkerboards, drifting gratings, white noise and sparse noise. (C) Representative 5 second trace of behavioral data acquired during free movement including locomotor speed, head velocity, eye velocity and gaze velocity (sum of head and eye velocity). Gaze shift events in either direction are marked with red triangles. Horizontal line in speed trace denotes threshold (2cm/s) for stationary vs locomoting. (D) Scatterplot of head and eye velocities for example recording in C, including gaze shifting (red), compensatory (black) and intermediate movements (grey). (E) Gaze shift movements. (*Left*) Schematic of head and eye movements that generate a gaze shift movement. (*Center Left*) Average eye, head and gaze velocities of all gaze shift movements before DOI administration. (*Center Right*) Average eye, head and gaze velocities of all gaze shift movements after DOI administration. (*Right*) Average ± s.e.m. change in the number of gaze shift movements evoked by administration of DOI (n = 10) or saline (n = 4). Calculated as the number of events after injection minus the number of events before injection. Black dot is the example recording used in C and D.

During free movement, animals coordinate their head and eye movements to sample the visual scene in a “saccade-and-fixate” pattern, which generates periods of stabilized retinal input during locomotion (Figure 1C). This pattern of visual processing occurs across the animal kingdom^18^ and under a variety of behavioral contexts^15,16,19,20^. While primates can make targeted saccades without moving their head, in mice, eye movements are generally coupled with a head movement. Therefore, during a head movement, mice can either move their eyes in the opposing direction to counteract the head movement, stabilizing the gaze, or move abruptly in the same direction as the head movement, resulting in a change in retinal input; we refer to these movements as compensatory or gaze-shifting, respectively^15,16,21^. The categorization of these two movements can be visualized by plotting eye vs head velocity (Figure 1D). Compensatory movements fall along the diagonal, indicating equal and opposite head and eye movements, whereas gaze-shifting movements extend orthogonal to the diagonal, representing eye and head movements in the same direction. Here we limited our investigation to horizontal head/eye movements because in mice vertical head/eye movements are largely compensatory^15,21^. We define gaze-shifting and compensatory movements as previously outlined^16,22^ (see Methods). Here we focus our analyses on gaze-shifting head and eye movements as those result in new retinal input to the visual system.

### DOI alters the frequency of visual active sampling behaviors

Psychedelics, including DOI, alter many aspects of mouse behavior, including locomotor activity^8^, head twitch responses^7^ and forepaw stereotypy frequency^9^. DOI has also been shown to alter mouse behavior during an olfactory-guided search task, increasing olfactory active sampling behaviors while simultaneously reducing search accuracy^23^. Therefore, we first asked if DOI (n = 10) or saline (n = 4) altered the active sampling behaviors (i.e. gaze shifts) of free-moving mice during naturalistic vision. The average visuhead and eye velocities generated during gaze shifting movements before and after DOI administration were similar (Figure 1E), illustrating that DOI is not altering the way in which animals move during gaze shift events. DOI did alter the frequency of gaze shifts, increasing the average number of gaze shifts per minute by 5.56 ± 3.79, however this effect was highly variable across mice. The effect of saline injections was the opposite, decreasing the number of gaze shifts per minute by an average of 9.92 ± 4.89. The difference in saline and DOI’s effect on gaze shift frequency was significant (Figure 1E; Wilcoxon rank-sum test; *Z* = 1.98, p = 0.04). Interestingly, the effect of DOI on gaze shift frequency was highly correlated to its effect on locomotion (Figure S1B; Pearson’s correlation coefficient; r = 0.89, p = 5.2e-4), suggesting DOI is increasing animal engagement in the arena resulting in increased active sampling behaviors.

### DOI attenuates visually-evoked activity during active sampling

Previous work has demonstrated that DOI attenuates overall visually-evoked responses in V1 during head-fixation^11^. Because gaze shifting movements initiate a sequence of visual activity in V1 evoked by the change in retinal input^16^ we asked if DOI altered the visually-evoked activity initiated by gaze shifts in freely moving mice. We recorded visual activity from V1 of 12 mice before and after subcutaneous administration of either DOI (n = 8, 418 cells) or a saline control (n = 4, 142 cells). DOI significantly decreased the percentage of neurons in our recordings that were considered responsive to gaze shifts (a change in the average gaze shift evoked absolute firing rate of >1 spike per second and >3.5 standard deviations above the baseline firing rate) from an average of 83.3% ± 2.1% before DOI to 71.9% ± 2.2% after DOI (Figure 2A; Wilcoxon signed-rank test; *Z* = 2.73, p = 0.003). This was not the case with saline control injections (Figure 2A; Wilcoxon signed-rank test; before saline: 88.6% ± 5.1%, after saline: 82.3% ± 6.1%; *Z* = 0.72, p = 0.24).

**Figure 2.**
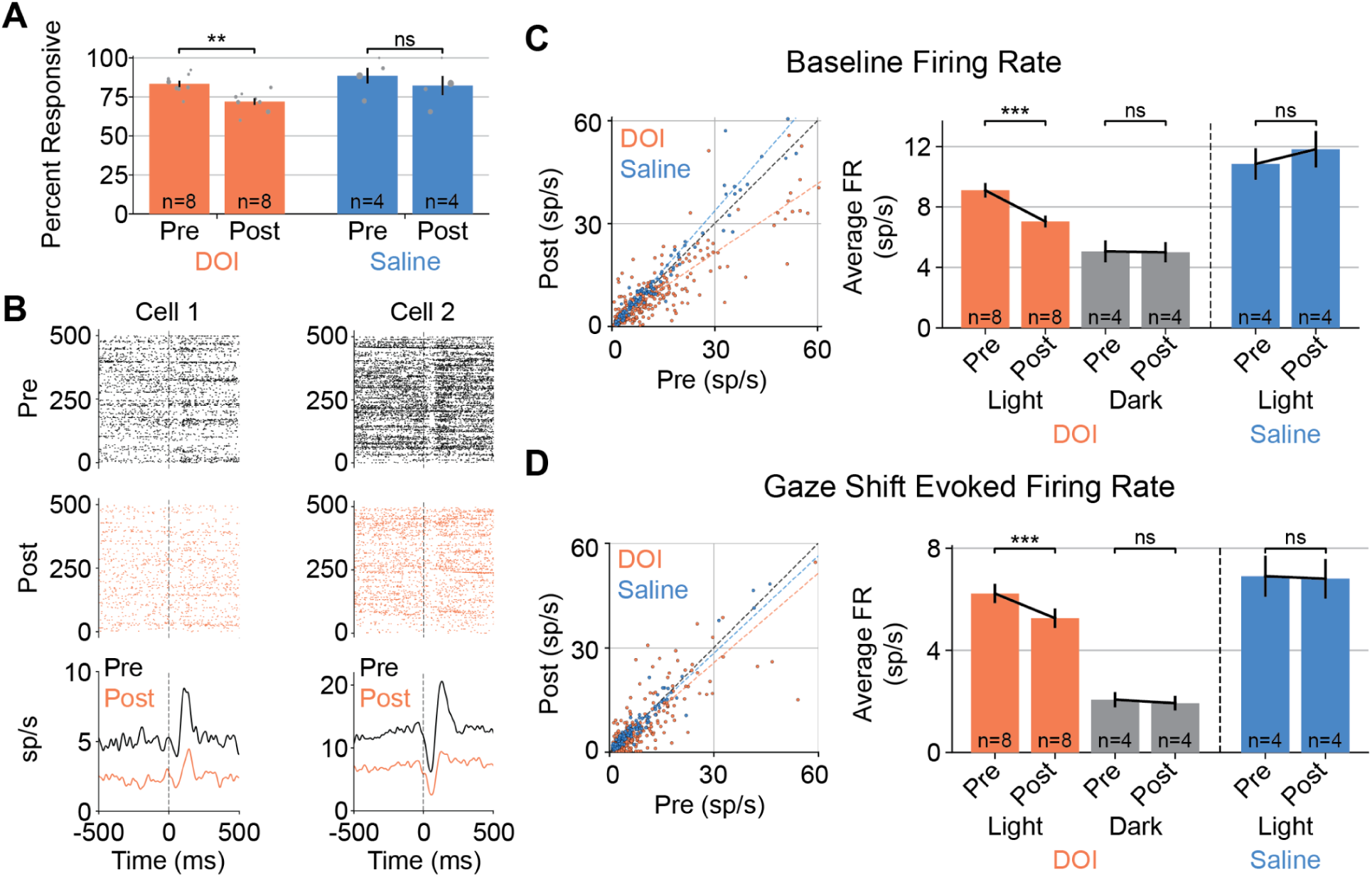
DOI suppresses visually-evoked responses in V1 during free movement. (A) Percentage of single units in each recording that were responsive to gaze shifts before and after administration of DOI or saline. Each dot represents a single recording and the size of each dot represents the percentage of the total pseudopopulation coming from that recording. (B) Example spike rasters to gaze shift responses before (*top*) and after (*middle*) DOI administration as well as their corresponding PETHs (*bottom*). Dotted line indicates onset of gaze shifts. (C) Baseline firing rates before and after DOI or saline administration. (*Left*) Scatterplot of baseline firing rates before and after DOI or saline. Black dotted line denotes y = x. Orange and blue dotted lines are linearly fit to the DOI and saline datasets, respectively. (*Right*) Average ± s.e.m. baseline firing rate across the DOI and saline populations before and after injections in the light. In a subpopulation of DOI mice, we recorded before and after injection in complete darkness. (D) Same as D but for the gaze shift evoked firing rate.

On average, DOI had a net suppressive effect on V1 neurons’ gaze shift responses, as previously observed for other sensory responses. Figure 2B shows two representative gaze shift responsive neurons before and after administration of DOI that illustrate the two main effects DOI had on gaze shift responses. First, DOI substantially suppresses the baseline firing rate (average firing rate from 800ms to 250ms before gaze onset). Importantly, during free movement this baseline period consists of ongoing visual processing outside of gaze shifts, rather than the absence of visual input as in a traditional pre-stimulus interval. Across the entire population, DOI significantly attenuated the baseline firing rate of gaze responsive neurons from an average of 9.16 ± 0.54 spikes per second before to 6.84 ± 0.43 spikes per second after (Figure 2C; Wilcoxon signed-rank test; *Z* = 4.35, p = 6.80e-6). Saline injections did not have a significant effect on the baseline firing rate of gaze responsive neurons (Figure 2C; Wilcoxon signed-rank test; *Z* = 0.35, p = 0.64). Second, DOI attenuated the gaze shift evoked firing rate (maximum firing rate from -50ms to 350ms after gaze onset after subtracting baseline). The average gaze shift evoked response across the population decreased from 6.22 ± 0.44 spikes per second before DOI to 5.30 ± 0.43 spikes per second after DOI (Figure 2D; Wilcoxon signed-rank test; *Z* = 3.94, p = 4.02e-5); whereas saline injections did not significantly affect the gaze shift evoked firing rate (Figure 2D; Wilcoxon signed-rank test; *Z* = 0.04, p = 0.49). In a subset of mice (n = 4, 134 cells), we recorded in complete darkness before and after DOI administration and found no significant effect of DOI on either the baseline (Figure 2C; Wilcoxon signed-rank test; Z = 0.20, p = 0.42) or gaze shift evoked firing rate (Figure 2D; Wilcoxon signed-rank test; Z = 1.39, p = 0.08), highlighting that DOI is attenuating visually-evoked activity in V1 during free movement.

### DOI alters gaze shift response magnitudes in temporally dependent manner

While the net effect of DOI across the population was suppressive, its effects on individual cell’s responses was diverse and we observed both facilitation and suppression (Figure 2C, D). We asked if these diverse effects could be explained by differences in gaze shift response dynamics. Gaze shifts evoke a diversity of visual responses in V1 that vary in their latency and polarity^16^. These can be broadly clustered into four main groups: early positive, late positive, biphasic and negative responses (Figure 3A), that collectively generate a continuous sequence of activity. To test if DOI affected these gaze shift clusters differently we clustered our datasets identically to Parker et al., 2023, based on each neuron’s pre-injection response. The distribution of early (DOI: 68 cells, saline: 27 cells), late (DOI: 68 cells, saline: 17 cells), biphasic (DOI: 84 cells, saline: 22 cells), negative (DOI: 49 cells, saline: 26 cells) and unresponsive clusters (DOI: 149 cells, saline: 50 cells) in our datasets were not significantly different from each other (Figure S2A; Chi-square test; X^2^_(4)_ = 6.39, p = 0.17). Additionally, the average cluster responses were similar before DOI or saline (Figure S2B), collectively illustrating similar pre-injection datasets. DOI attenuated the average gaze shift responses of all four clusters and this was significantly different than the effect of saline in all but the early cluster (Figure 3A).

**Figure 3.**
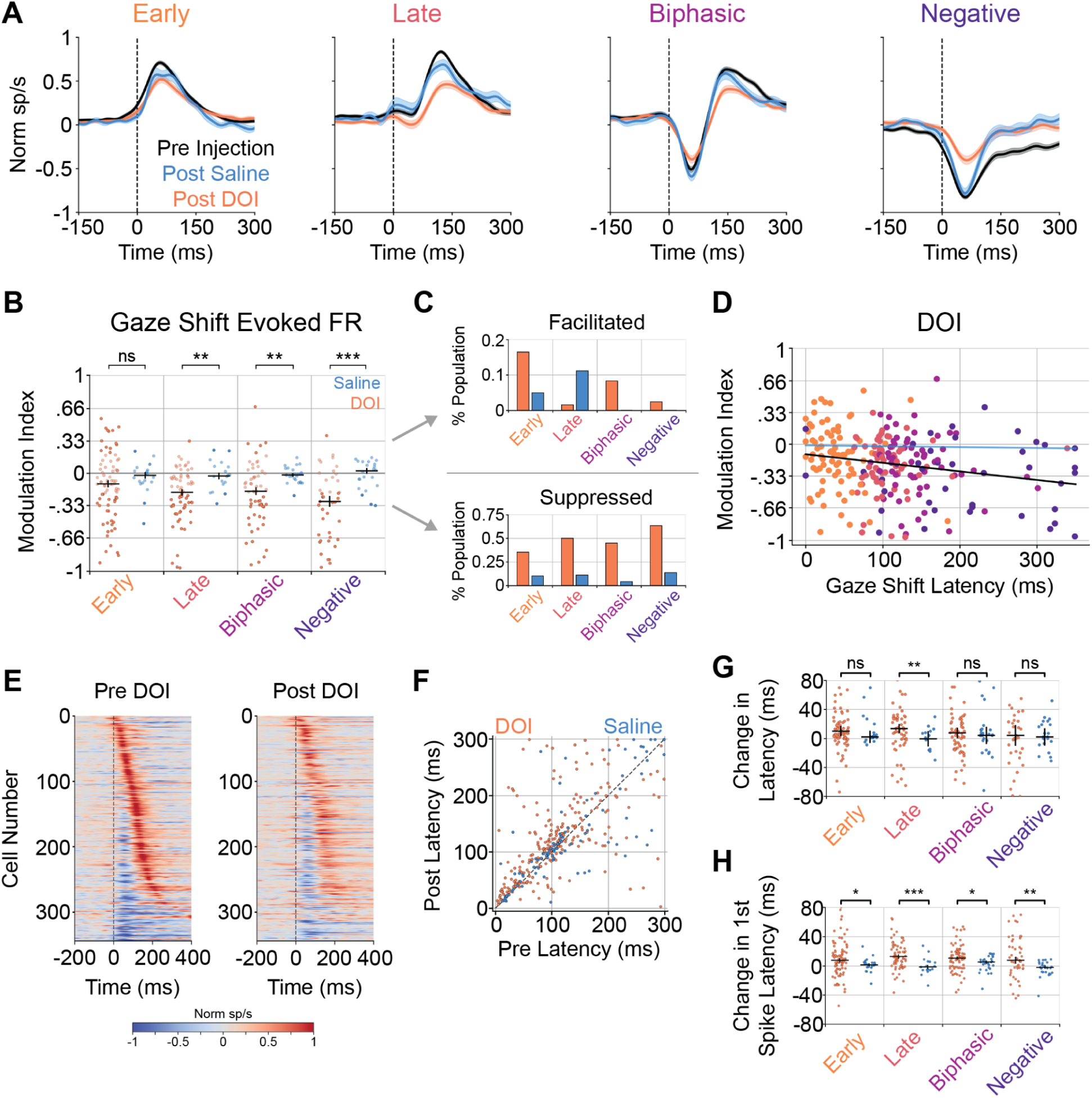
DOI alters gaze shift responses in cluster specific manner. (A) Average ± s.e.m. gaze shift response PETHs before or after DOI or saline injections clustered as in Parker et al., 2023. Cluster identity was determined using the pre DOI gaze shift responses. Average PETHs were similar before DOI or saline injection and were combined in these plots. (B) Modulation index of effect of DOI or saline administration on gaze shift evoked responses sorted by gaze shift cluster identity. Positive or negative values indicate injection-induced facilitation or suppression of gaze shift evoked responses, respectively. Median ± s.e.m. values included for each condition shown in black. Values greater than 0.2 or less than -0.2 were considered substantially modulated by the injection. Cells that were not substantially modulated were plotted as slightly transparent. (C) Percentage of each cluster that had modulation indices greater than 0.2 (facilitated) or less than -0.2 (suppressed) for DOI and saline datasets. (D) Gaze shift evoked firing rate modulation index vs gaze shift response latency for the DOI population. Black line is linear fit to the DOI data, blue line is linear fit to the saline data (not plotted, see Figure S2C). (E) Temporal sequence of normalized gaze shift responses sorted along the y-axis by response latency before (*left*) and after (*right*) DOI. In both plots cells are sorted by their pre-injection response latencies. Dotted line denotes gaze shift onset. (F) Scatterplot comparing gaze shift latencies before and after DOI or saline administration. (G) Change in gaze shift latencies induced by injection sorted by gaze shift response cluster. Positive and negative values indicate injection-induced increases and decreases in response latency, respectively. Median ± s.e.m. values included for each condition. (H) Same as G but for first spike latency.

To quantify the effect of DOI administration on individual cell gaze shift responses, we calculated a modulation index (Figure 3B, see methods). Values less than zero reflect suppression of gaze shift responses while values greater than zero reflect facilitation. This revealed a striking trend across gaze shift clusters, as the effect of DOI, but not saline, became more consistently suppressive as a function of gaze shift cluster latency (Figure 3B). While the effect of DOI on the early cluster was heterogeneous, generating both facilitation and suppression of visual responses, its effect on the negative cluster was almost entirely suppressive. By setting a modulation threshold of >0.2 or <-0.2, we quantified the percentage of neurons in a cluster whose gaze shift responses were considerably facilitated or suppressed by either DOI or saline, respectively (Figure 3C). The percentage of cells in the early, late, biphasic and negative clusters that were facilitated by DOI were 17%, 2%, 8% and 3%; respectively. Conversely, the proportion of cells in each cluster that were suppressed by DOI were 35%, 50%, 45% and 64%, respectively. Thus, the effect of DOI on gaze shift responses during free movement depends on the latency of a cell’s response, becoming more consistently suppressive as a function of gaze shift response latency. To highlight this effect, we plotted each cell’s gaze shift modulation index against its gaze shift response latency for our DOI (Figure 3D) and saline datasets (Figure S2C). Across the population there was a significant negative relationship between gaze shift modulation index and gaze shift response latency for DOI (Pearson correlation coefficient; r = -0.22, p = 6.5e-4) but not saline (Pearson correlation coefficient; r = -0.06, p = 0.60). A similar trend was present in the effect of DOI on the baseline firing rate; a cell’s gaze shift response latency was predictive of the direction and magnitude of DOI’s effect on the baseline firing rate (Figure S3A-C). Finally, comparing the effect of DOI on the gaze shift response to its effect on the baseline firing rate revealed a significant positive relationship (Figure S3D; Pearson correlation coefficient, r = 0.82, p = 5.60e-105), indicating that DOI had similar effects on a cell’s visual response both during and between gaze shifts.

### DOI doesn’t alter coarse-to-fine pattern between gaze shift response latency and SF preference

This latency-specific effect of DOI on gaze shift responses is particularly interesting given that these responses form a temporal sequence of activity that encodes visual information in a “coarse-to-fine” pattern^16^, wherein lower spatial frequency (SF) components of the visual scene are encoded prior to higher SF details^24,25^. Thus, we next sought to investigate if DOI affected this coarse-to-fine pattern of visual activity, by disrupting the relationship between response latency and SF-preferences of gaze shift responsive cells.

Plotting the PETH of all responsive cells, sorted by the latency of their peak positive gaze shift response before DOI or saline administration, revealed a dynamic sequence of activation to gaze shifts before DOI (Figure 3E) or saline (Figure S2D), similar to that described previously ^16^. We used cross-validation to confirm the temporal pattern (Figure S4; Pearson’s correlation coefficient; DOI: r = 0.67, p = 3.43e-46; saline: r = 0.58, p = 1.04e-11). Sorting the post injection responses by their pre injection latencies highlighted that the temporal pattern of activity following a gaze shift is not qualitatively different after DOI (Figure 3E) or saline (Figure S2D). Furthermore, there was a significant positive relationship between the pre and post injection latencies for both populations (Figure 3F; Pearson’s correlation coefficient; DOI: r = 0.61 p = 1.32e-26; saline: r = 0.81, p = 7.42e-24). While DOI did subtly increase the median gaze shift latency of all clusters, this was only significantly different than the effect of saline in the late cluster (Figure 3G; Wilcoxon rank-sum test; p = 0.004). Interestingly, DOI did significantly increased the median first spike latency of all clusters relative to saline (Figure 3H; Wilcoxon rank-sum test; early: p = 0.02, late: p = 6.3e-5, biphasic: p = 0.04, negative: p = 0.001).

We next asked if DOI altered the SF tuning of gaze shift responsive neurons by presenting the animal (DOI: n = 8; saline n = 4) with drifting sinusoidal gratings at three SFs (0.02, 0.08, 0.32 cycles per degree, cpd) during the head-fixed portion of our recordings (Figure 1B). Consistent with previous reports^11^, DOI had a suppressive effect on the average response to all grating stimuli (Figure 4A). For each cell responsive to the grating stimuli (DOI: 190 cells, saline: 87 cells) we calculated a modulation index using the maximum evoked response to each SF before and after either DOI or saline administration. Across the population, DOI suppressed the median maximum evoked response to each of the three tested SFs, however this was only significantly different than the effect of saline for the highest tested SF (Figure 4B; Wilcoxon rank-sum test; 0.02 cpd: p = 0.09, 0.08 cpd: p = 0.54, 0.32 cpd: p = 8.6e-3). Next, we calculated the weighted SF preference of each gaze responsive cell before and after injection of DOI or saline and found a significant positive relationship between the pre and post weighted SF preferences in both populations (Figure 4C,E; Pearson’s correlation coefficient; DOI: r = 0.82, p = 1.04e-41; saline: r = 0.95, p = 5.65e-41). Plotting the weighted SF preferences by gaze response cluster before DOI (Figure 4D) or saline (Figure 4F) demonstrated a clear coarse-to-fine progression with early, late, biphasic and negative cells preferring progressively higher SFs. Neither DOI nor saline significantly affected the median SF preferences of these gaze response clusters with the exception of DOI subtly but significantly reducing the median SF preference of negative cells (Figure 4F; Wilcoxon signed-rank test; p = 0.01). Thus, while DOI administration caused drastic changes in gaze shift response amplitudes, it did not substantially disrupt the coarse-to-fine relationship between response latency and SF preference across the population response.

**Figure 4.**
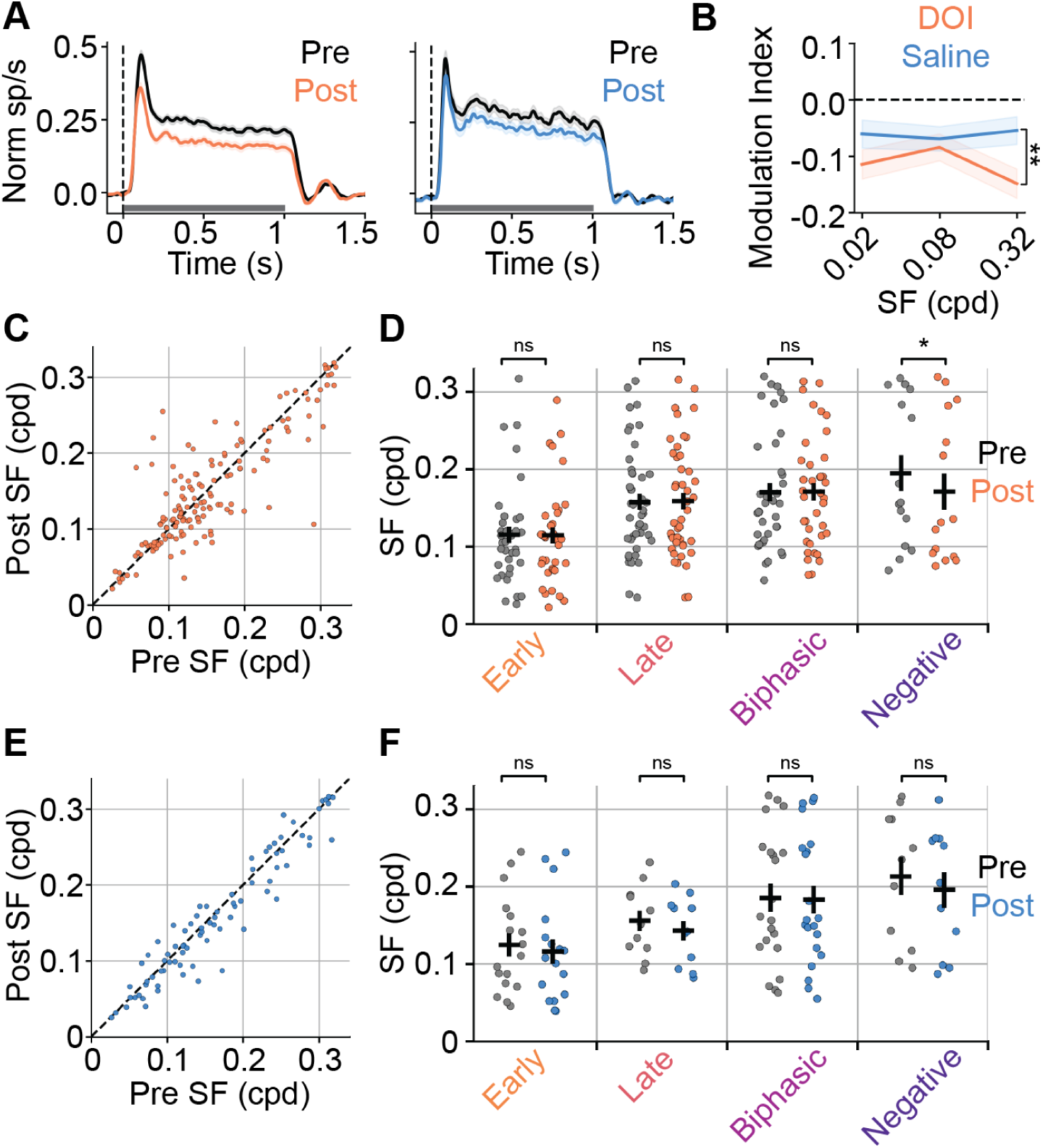
DOI doesn’t alter SF preferences of gaze shift responsive neurons. (A) Average ± s.e.m. response to all presented grating stimuli before or after injection of DOI (left) or saline (right). Dotted vertical line denotes stimulus onset, grey horizontal line indicates stimulus duration. (B) Median modulation index across DOI and saline datasets for each tested SF. Maximum evoked responses were used to calculate the modulation index. (C) Scatterplot comparing preferred SF before and after DOI administration. The dotted line indicates y = x. SF preferences were calculated as a weighted mean of responses. (D) Preferred SFs before and after DOI administration sorted by gaze shift response cluster. Median ± s.e.m. are shown for each condition.

## Discussion

To determine the impact of psychedelics on active visual processing under naturalistic conditions, we recorded the activity of V1 neurons in freely moving mice before and after administration of the psychedelic compound DOI. Consistent with previous studies we observed a net suppressive effect of DOI on visually-evoked activity in V1^11,12^, however DOI’s effect on individual neurons varied dramatically. We found a striking relationship between the direction of DOI’s modulation on gaze shift responses and the latency of those responses. Short latency cells displayed DOI-induced facilitation and suppression whereas the effect of DOI became more consistently suppressive in gaze shift cells with longer response latencies. Thus, examining the impact of a psychedelic in the context of natural, active vision revealed a more complex and dynamic view of its impact on neural coding.

Our findings complement previous work studying the effect of psychedelics on visual function in humans^12^ and head-fixed mice^11^; building upon these studies by providing important naturalistic context to our understanding of the diverse effects that psychedelics have on sensory processing. Importantly, these findings suggest potential mechanisms for the altered visual perception induced by psychedelics. The diverse modulatory effects DOI has on visual responses, simultaneously facilitating and suppressing different neural populations, likely disrupts the decodability of visual representations. Unlike a uniform gain change, which preserves the relative activity between neurons and the structure of the population code, a heterogeneous gain modulation, as we observe with DOI, distorts these activity patterns in a way that could disrupt decodability of the population code. Consistent with the idea that heterogeneous modulations can be particularly detrimental to neural representations, work in macaque V1 has demonstrated that stimulus uncertainty is represented as increases in cross-neuron gain variability^26^.

The influence of this diverse modulation on visual encoding is especially notable in the context of the coarse-to-fine processing that unfolds following gaze shifts. A majority of facilitated cells were from the population that responds at low latencies and prefer low SFs, whereas cells that respond at longer latencies and prefer higher SFs, were most consistently suppressed. According to Marr’s primal sketch theory, the role of early visual processing (i.e., coarse) is to establish the broad structural scaffold of a scene while later processing resolves its finer details^27^. Disruptions to the relative balance between these coarse and fine representations, like we observe here with DOI, could therefore impair the visual system’s ability to appropriately structure and subsequently fill in the visual scene.

This imbalance across the temporal sequence may also be understood within a broader framework linking psychedelic hallucinations to disruptions in top-down and bottom-up processing across the visual hierarchy^11,28–30^. It is now well established that psychedelics have a net suppressive effect on bottom-up visually-evoked activity^11,12^, and our finding that this suppression is particularly strong in cells that encode the high-SF details of a visual scene suggest top-down predictive signals might disproportionately contribute to the representation of these high-SF details. Thus, our data support a framework in which visual hallucinations result from an imbalance in coarse and fine input integration along with an over reliance of top-down predictive inputs in the representation of high-SF details of a scene, consistent with the frequent appearance of textures, geometric patterns, and other “form constants” in psychedelic imagery^31,32^. Future studies integrating sensory processing under psychedelics with behavioral^33^ or computational^34^ readouts of perception will be promising research directions to empirically test this framework and more directly link the neural and perceptual consequences of psychedelics.

Our findings also raise a number of interesting questions regarding the neuronal and circuit mechanisms of DOI’s effect on visual processing. First, why are some neuronal responses enhanced by DOI while others are suppressed? Clustering spike waveforms into narrow and wide spiking neurons, representing putative parvalbumin expressing (PV+) interneurons and excitatory neurons, respectively, failed to explain our diverse bidirectional effects (Figure S5A-C). However, it is possible that other cell types, defined genetically or by their projection specific patterns^35,36^ might underlie these diverse modulatory effects of DOI. Alternatively, these effects might relate to laminar differences in serotonergic receptor expression patterns^37^. Consistent with this, we found that cells that were facilitated by DOI were more likely to be near layer 5 (L5) of V1 than suppressed cells (Figure S5D). L5 of the neocortex is particularly enriched with 5HT-2A receptors^37^ and recent work has shown that serotonergic agonists like DOI directly excite L5 neurons in the prefrontal cortex^38^. Thus, one compelling possibility is that the DOI-induced facilitation of gaze shift responses results from direct agonism of these L5 receptors while suppression reflects activation of other serotonergic receptor populations or network level effects.

Second, what are the neuronal and circuit mechanisms that lead to a progressive increase in DOI-induced suppression following a gaze shift? This effect is likely related to the circuit mechanisms that shape the coarse-to-fine temporal sequence following a gaze shift. One possibility is distinct visual inputs with different spatial and temporal tuning (i.e., magno- and parvo-cellular pathways) contributing differently throughout the temporal response, with DOI differentially affecting these pathways. While there is some evidence for these distinct pathways in rodents, or at least a continuum between the two^39,40^, the impact of serotonergic psychedelics on these pathways has not yet been studied. Alternatively, it is intriguing to hypothesize that a recurrent circuit mechanism shapes the coarse-to-fine temporal structure following a gaze shift and this recurrency progressively amplifies DOI’s suppressive effect across the duration of the sequence, thereby resulting in a shift towards net suppression and disruption of the coarse-to-fine balance.

The findings presented here underscore the importance of studying sensory processing in naturalistic contexts, as our work reveals a richer and more nuanced view of the impact of psychedelics than previously possible with constrained paradigms. By recording from freely moving animals engaged in natural exploration and experiencing complex real-world visual inputs, we uncovered a diverse and temporally dynamic modulation of sensory activity that is linked to active sampling behaviors. Identifying the specific neuronal and circuit-level mechanisms that drive these effects will lead to a deeper, more empirical understanding of both normal sensory function and the profound alterations in perception that characterize the psychedelic state. Ultimately, bridging the gap between large-scale neural dynamics and perceptual readouts offers a promising path toward unlocking how the brain constructs a coherent visual world and how that construction can be so strikingly transformed.

## Author Contributions

RJS and CMN conceived the project. CMN supervised all aspects of the project. RJS and CWF collected all the data from mouse experiments. RJS led data analysis with contributions from DMM. RJS, CWF and CMN contributed to writing and editing the manuscript.

## Acknowledgements

We thank Dr. Angie Michaiel, Dr. James Murray, Dr. Phil Parker, and members of the Niell Lab for feedback on previous drafts of the manuscript. This work was supported by NIH grants RF1NS127305 (CMN), R01DA055439 (CMN), R25MH131653 (CMN, RJS) and F32EY032360 (RJS).

## Methods

### Animals

All procedures were conducted in accordance with National Institutes of Health guidelines and were approved by the University of Oregon Institutional Animal Care and Use Committee. 4-12 month-old mice (Mus musculus, C57BL/6J, Jackson Laboratories and bred-in house) were kept on a 12-h light/dark cycle. In total, 10 male and 6 female mice were used for this study. Mice were housed with sibling cage mates until the beginning of experiments and then they were singly housed. Humidity was between 40-60% and temperature was 21±1°C. Data collection and analysis were not performed blind due to the condition of the experiments.

### Drug Dosage and Administration

DOI (2,5-dimethoxy-4-iodoamphetamine; Sigma, 10mg/kg in saline) or saline (0.9% NaCl) were administered subcutaneously halfway throughout the recording using previously published protocols from studies investigating the effects of DOI on sensory processing^10,11^. Subcutaneous injection was used rather than intraperitoneal to prevent having to remove the animal from the head-fixed setup and minimize animal handling and electrode movement during the recording. We intentionally chose the concentration of 10 mg/kg of DOI to be at the high end of standards in the literature, which range from 1-10 mg/kg intraperitoneal^7,10,11^; to account for differences in pharmacokinetics between these methods of administration^41^. There was no blinding regarding the reagent used.

### Surgery and Habituation

Mice were initially implanted with a titanium headplate over V1 to allow for head-fixation and attachment of head-mounted experimental hardware. After 3 days of recovery, widefield imaging was performed to help target the electrophysiology implant to the approximate center of the left monocular V1. A miniature connector (Mill-Max 853-93-100-10-001000) was secured to the headplate to allow repeated, reversible attachment of a camera arm, eye/world cameras and inertial measurement unit (IMU)^15,16,42^. To simulate the weight of the real electrophysiology drive for habituation, a ‘dummy’ electrophysiology drive was glued to the headplate. Animals were handled by the experimenter for several days before surgical procedures, and subsequently habituated (∼45 minutes total) to the spherical treadmill and freely moving arena with hardware tethering attached for several days before experiments.

The electrophysiological implant was performed once animals moved comfortably in the arena. A craniotomy was performed over V1, and a linear silicon probe (128 channels, Diagnostic Biochips P128-6 or P64-10-D) mounted in a custom three-dimensionally printed drive (Yuta Senzai, UCSF) was lowered into the brain using a stereotax to an approximate tip depth of ∼750 μm from the pial surface. The surface of the craniotomy was coated in artificial dura (Dow DOWSIL 3-4680) and the drive was secured to the headplate using light-curable dental acrylic (Unifast LC). A second craniotomy was performed over the right cerebellum and a reference wire was inserted into the brain. The opening of the reference wire craniotomy was covered with a small amount of ophthalmic ointment and the wire was glued in place with UV light cure glue (Loctite AA3972). Animals recovered overnight and experiments began the following day.

### Hardware and Recording

The camera arm was oriented approximately 90 deg to the right of the nose and included an eye-facing camera (iSecurity101 1000TVL NTSC, 30 frames/s (fps), interlaced), an infrared light-emitting diode to illuminate the eye (Chanzon, 3-mm diameter, 940-nm wavelength), a wide-angle camera oriented towards the mouse’s point of view (BETAFPC CO1, 30 fps interlaced) and an IMU acquiring three-axis gyroscope and accelerometer signals (Rosco Technologies; acquired at 30kHz, downsampled to 300 Hz and interpolated to camera data)(Figure 1A). Fine-gauge wire (Cooner, 36 AWG, no. CZ1174CLR) connected the IMU to its acquisition box, and each of the cameras to a USB video capture device (Pinnacle Dazzle or StarTech USB3HDCAP). A top-down camera (FLIR Blackfly USB3, 60fps) recorded the mouse in the arena.

The electrophysiology headstage (built into the silicon probe package) was connected to an Open Ephys acquisition system via an ultra-thin cable (Intan no. C3216). Electrophysiology data were acquired at 30 kHz and bandpass filtered between 0.01 Hz and 7.5 kHz. We first used the Open Ephys GUI (https://github.com/open-ephys/plugin-GUI) to assess the quality of the electrophysiology data, then recordings were performed in Bonsai using custom workflows (https://github.com/nielllab/FreelyMovingEphys). System timestamps were collected for all hardware devices and later used to align data streams through interpolation.

During experiments, animals were first placed in the freely-moving arena which they could explore for 45 minutes and then subsequently head-fixed on a spherical treadmill for approximately 30 minutes to permit measurement of visual tuning properties using traditional methods (Figure 1B). Animals were then given a subcutaneous injection of either saline (0.9% NaCl) or DOI (2,5-dimethoxy-4-iodoamphetamine; Sigma, 10mg/kg in saline) and mice were monitored for front paw stereotypy^9,11^, which DOI reliably induced within 5-7 minutes following the injection. After waiting 10 minutes, experiments were repeated in reverse order to minimize the number of times the animal had to be moved from the head-fixed preparation to the freely-moving arena (Figure 1B). In total, recordings took ∼2-3 hours.

For head-fixed experiments, a 70-cm monitor (BenQ GW2780) was placed approximately 27.5 cm from the mouse’s right eye, and visual stimuli were presented using Psychtoolbox-3^43^. Head-fixed stimuli were also recorded using the head-mounted world camera. First, we presented 15 min of a band-limited Gaussian noise stimulus^39^ (SF spectrum 0.05 cpd to 0.12 cpd, flat TF spectrum with a low-pass cutoff at 4 Hz) to confirm V1 targeting based on spike-triggered average receptive fields. Next, we presented a contrast-reversing square-wave checkerboard stimulus for 5 mins with a SF of 0.04 cpd and TF of 2 Hz. Drifting sinusoidal gratings were presented at eight evenly spaced directions of motion for three SFs (0.02, 0.08, 0.32 cpd) and two TFs (2, 8 cps) for 15 min, with a 1-s stimulus duration and 0.5-s gray ISI, and stimulus conditions randomly interleaved (total of 12 presentations per stimulus).

The arena was approximately 48-cm long by 37-cm wide by 30-cm high (Figure S1A). A 61-cm monitor (BenQ GW2480) covered one wall of the arena, while the other three walls were clear acrylic covering custom wallpaper including black and white high-SF and low-SF gratings and white noise. During recordings, the arena’s stimulus monitor played a sparse moving noise stimulus containing full- and minimum-luminance spots 4, 8 and 16 deg in diameter. Each spot was assigned to move in one of eight evenly spaced directions and at one of five speeds (10, 20, 40, 80, 160 deg per s). Spots appeared on the appropriate edge of the screen and moved across until they disappeared on the far edge. The floor was a gray silicone mat (Gartful) and was covered with black and white Lego bricks to provide three-dimensional visual contrast. Small pieces of tortilla chips (Juanita’s) were lightly scattered around the arena to encourage foraging during the recording; however, animals were not water or food restricted.

For dark recordings the entire experimental enclosure was sealed in light-blocking material, and all potential light sources within the enclosure and all external light sources were turned off, to ensure complete darkness. Prior to darkness experiments one drop of 2% pilocarpine HCl ophthalmic solution was applied to the animal’s right eye to constrict the pupil and allow for accurate eye tracking as described in ^16^. Once the pupil was restricted enough for tracking in the dark (3-5 min) the animal was moved from the spherical treadmill to the dark arena for the recording. The recording lasted approximately 30 min or until the effect of pilocarpine wore off followed immediately by the light recording.

### Data Preprocessing

Electrophysiology data was preprocessed following the same methodology as ^16^. Raw electrophysiology data from head-fixed and free moving recordings in the same session were concatenated into a single file for spike sorting, allowing us to track single units across the entire experiment. Common-mode noise was removed by subtracting the median across all channels at each timepoint. Spike sorting was performed using Kilosort 4 (https://github.com/MouseLand/Kilosort), and single units were isolated and selected using Phy 2.0 (Phy 2.0 beta 5; https://github.com/cortex-lab/phy) based on numerous parameters: contamination index (< 30%), firing rate (mean > 0.5 Hz across entire recording), autocorrelogram, and waveform shape. Following spike sorting the data were then split back out into individual stimuli recordings for further analysis.

To group units into narrow- and broad-spiking putative cell types, we first normalized the mean spike waveform of each unit using the equation below, where W is the waveform and Wb is the mean of the first 200 μs. We performed k-means clustering on the normalized waveforms (k = 2). The resulting clusters closely resembled the segregation of narrow- and broad-spiking units in previous studies based on explicit waveform criteria^44^.

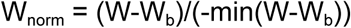

Laminar depth was calculated from the multi-unit local field potential (LFP) of head-fixed neural data. The mean-centered LFP of each channel for a given shank of the linear silicon probe was bandpass filtered (1 to 300 Hz) and the power for the channel was calculated. The position of the maximum LFP power along the ordered sites of each shank was considered the center of layer 5 of the cortex^45^.

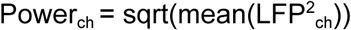

We extracted pupil position from the eye camera data^15^. Briefly, eye camera data were first deinterlaced to achieve 60-fps video, then eight points around the pupil were tracked with DeepLabCut^46,47^. We then fit an ellipse to these eight points and computed pupil position in terms of angular rotation. The head-mounted world camera was deinterlaced to achieve 60-fps video and distortions from the camera lens were corrected using OpenCV. The mouse’s position in the arena was tracked with DeepLabCut using top-down camera frames. We defined running as periods when the animal’s neck point had a velocity of >2 cm/sec and a stationary state as periods when the neck point had a velocity of <2 cm/sec (Figure 1C).

Horizontal head rotation velocity was extracted from the IMU, converted to deg/sec and interpolated to eye camera frame timestamps. We defined leftward and rightward directions as the direction of the head movement from the animal’s perspective. For example, a movement of the right eye in the nasal direction would be a leftward movement. To select eye/head movement onsets, we first determined if the head velocity was >60 deg/sec in the leftward or rightward direction. If a sufficiently large head movement was made, we then separated it into either gaze-shifting or compensatory movements using gaze velocity, where gaze is defined as the sum of horizontal eye and head velocities^15,16^. If there was a high gaze velocity (>240 deg/sec) concurrent with the head movement, the movement was considered gaze-shifting. Those that resulted in low gaze velocity (<120 deg/sec) were considered compensatory. Eye/head movements that resulted in the intermediate gaze velocities (>120 deg/sec and <240 deg/sec) were excluded from our analysis to avoid contamination between the two categories of movements. For head/eye movements spanning multiple eye camera frames, only the first time point, representing the onset of the movement, was used to calculate the onset of the PETH. Compensatory movements which occurred 250 ms before or after a gaze-shifting movement were excluded to avoid contamination by gaze shifts.

To maintain internal consistency with eye cameras and obviate the need for a separate video stimulus synchronization signal, we determined stimulus onset in head-fixed recordings directly from the head-mounted camera. For the reversing checkerboard stimulus that updated every 500 ms, we identified frame transitions from the head-mounted camera video based on *k*-means clustering (*k* = 2) of the video into the two separate contrasts, and selected transitions between the clusters. For drifting sinusoidal gratings, we determined direction of motion by computing optic flow from the worldcam video, SF based on the mean gradient magnitude from paired Sobel operators and TF from the mean Fourier transform of each pixel over time.

### Analysis of Neural Responses

All analyses were performed in Python (v.3.8, python.org). Neural responses were calculated as a PETH from electrophysiology spike times using kernel density estimation with a gaussian kernel with bandwidth (standard deviation) of 10 ms, sampled at 1 ms intervals. The PETH included neural activity from -1000 ms to 1000 ms around the event onset for all stimuli/events (unless noted otherwise). PETHs were calculated separately for pre and post injection recording periods. A modulation index was calculated for PETHs as the peak of the firing rate (*R*) from the event onset at -50 ms until 500 ms after the event, minus the baseline before the event (*R_b_*), which was calculated as the mean of *R* from -900 ms to -250 ms around the event onset (unless noted otherwise).

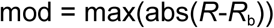

The modulation index from gaze-shift responses in the leftward and rightward directions were compared to determine a preferred direction. A direction selectivity index (DSI) was calculated as the difference between the preferred and nonpreferred maximum modulations divided by their sum.

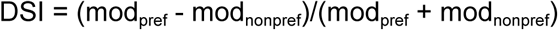

To normalize PETHs for each cell, we first subtracted the baseline *R_b_* from the PETH (*R*) to give an evoked firing rate and then divided by either the maximum of R before or after drug administration, whichever was larger, to maintain the effect of drug administration of the cell’s response throughout normalization. The maximum value of *R* was chosen during a response window (*R_rw_*) within a response range of -50 before to 500 ms after the event.

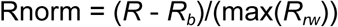

When normalizing the PETH of compensatory eye/head movements, we used that neuron’s *R_rw_*from gaze shifts in the preferred left/right direction and before or after drug administration, so that all eye/head responses were normalized relative to the neuron’s largest response.

For the drifting grating stimulus, the PETH was calculated similarly to other stimuli, but from -1500 ms before until 1500 ms after stimulus onset to capture the full stimulus and ISI interval. Normalization was also done similarly to other stimuli, where *R_b_* was calculated as the mean of *R* from -100 ms to -400ms around the stimulus onset.

Neurons were considered responsive to an eye/head movement onset if they changed their firing rate by at least 1 spike per second (sp/s) in the 350 ms following the onset of the event and if the absolute change in their firing rate during this 350 ms window rose to at least 3.5 standard deviations above *R_b_.* Neurons were considered responsive to onsets of head-fixed stimuli if they changed their firing rate by at least 10% and 1 spike per second (sp/s) in the 250 ms following the onset of the event. For the grating stimulus, suppressed-by-contrast cells were not considered responsive, and removed by requiring that grating PETHs have a peak firing rate >0.5 normalized sp/s during the 1 s of stimulus presentation.

The latency of a neuron’s peak response was calculated as the time point of the maximum firing rate of its normalized PETH in the period between 25 ms and 350 ms after the onset of the event. For temporal sequence plots, normalized PETHs were sorted by the latency of their peak for the preferred direction of gaze-shifting eye/head movements. Peak latency sorting was cross-validated by randomly assigning gaze-shifting events into a train set and a test set, calculating a PETH for each half of the data, and sorting the test set by the peak times of the training set (Figure S4).

We used a modulation index to quantify the effect of either DOI or saline administration on neural responses. We calculated the modulation index as the difference between the post and pre injection neural response divided by their sum. This resulted in a value between -1 and 1, where negative and positive values indicated injection-specific suppression and facilitation, respectively.

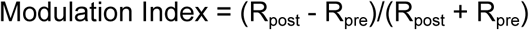

SF and TF preferences were based on the evoked firing rate for each stimulus condition, computed as the mean rate from 25 to 1000 ms following stimulus onset, minus the mean baseline rate in the -400 ms to -100ms before stimulus onset. The mean SF and TF tuning curves for all cells within gaze-shift clusters were calculated using each cell’s mean evoked firing rate for each SF/TF normalized by the maximum response to its best SF/TF, either before or after drug administration to preserve the effects of the drug on normalized responses.

Each cell’s weighted TF response, W_TF_, was determined using the mean evoked response, *R*, for each of the two presented TFs and at the cell’s preferred SF and orientation.

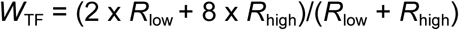

Likewise, a weighted SF response was calculated at the cell’s preferred TF and orientation using the mean evoked responses for each of the three presented SFs.

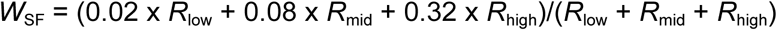

### Response Clusters

To cluster units with similar gaze-shift responses we followed methods laid out in ^16^. Briefly, we performed PCA on the normalized pre-injection gaze-shift response in each unit’s preferred direction in the period from -50 to 300 ms relative to the gaze-shift onset. Units that did not reach the above spike rate criteria had their input for PCA replaced with an array of zeros to exclude unresponsive PETHs from clustering, while seeding the formation of an unresponsive group of units from clustering. We used the first four principal components, which cumulatively explained 95% of the variance, to perform *k*-means clustering (*k* = 5), resulting in four clusters of responsive units and one cluster of unresponsive units.

### Statistical Analysis

Data were statistically analyzed using custom scripts written in Python (v.3.8, python.org). We used the Chi-square test, Wilcoxon rank-sum test, Wilcoxon signed-rank test, Kruskal-Wallis test and Kolmogorov-Smornov test to determine statistical significance. When appropriate we used one-tailed statistical tests. The Chi-square test was used for proportional comparisons. The Wilcoxon rank-sum test was used to compare two groups of independent samples. The Wilcoxon signed-rank test was used to compare paired samples. The Kruskal-Wallis test with post-hoc Dunn’s test was used to compare three or more independent samples. All error bars represent the standard error of the mean (SEM). In figures, * p < 0.05, **p <= 0.01, and ***p <= 0.001.

---

**Figure S1.**
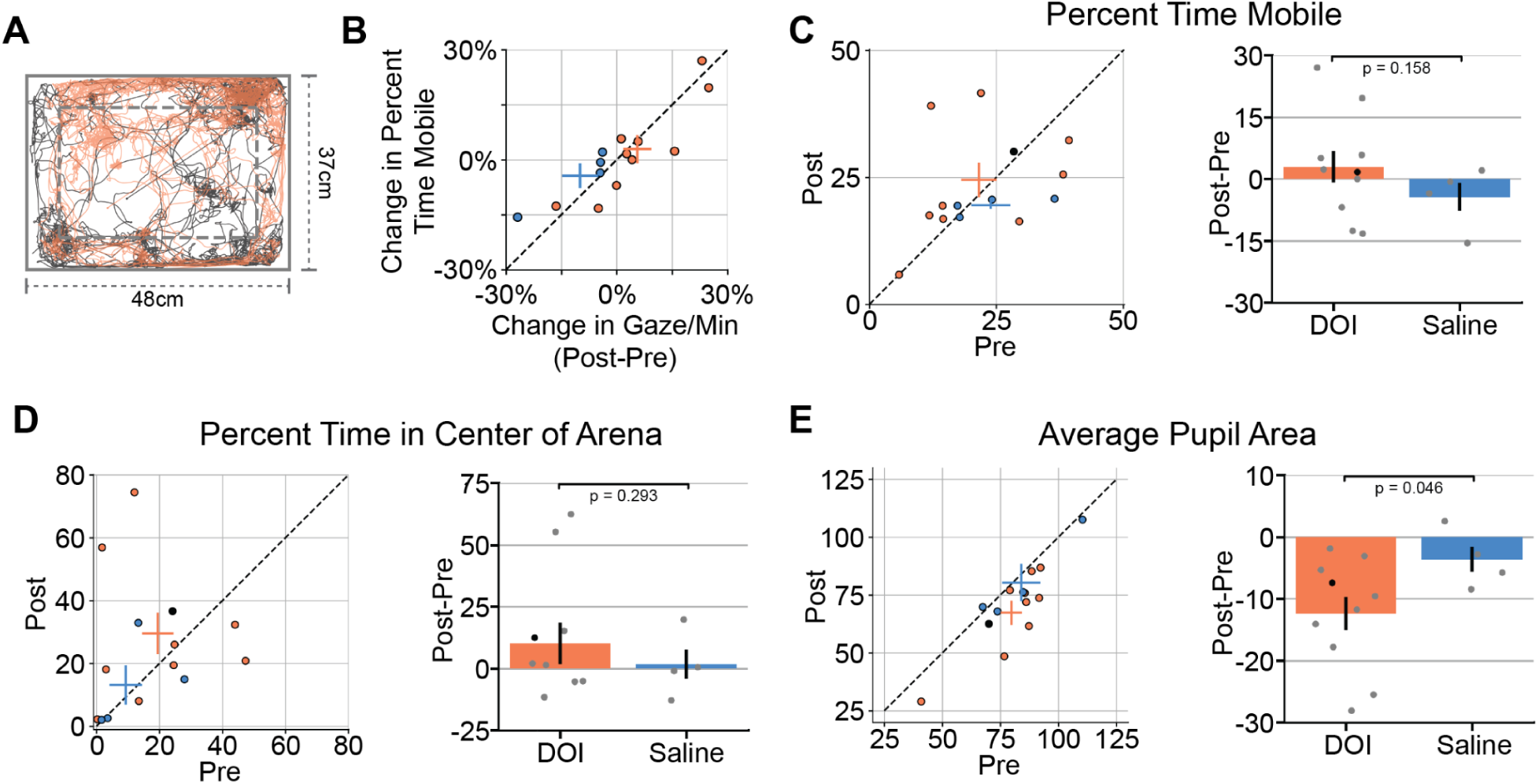
Effect of DOI on additional behavioral measures. (A) Top-down view of freely moving recording arena with example mouse position throughout recording period before and after DOI administration. Dotted inset box marks area used to quantify center and surround preferences of an animal. The arena is 37 cm wide and 48 cm long. Example mouse is the same as used in Fig 1. (B) Scatterplot comparing the change in gaze shift frequency and change in locomotor activity caused by administration of DOI (orange) or saline (blue). Mean ± s.e.m plotted for both populations. (C) Percentage of time mobile before and after DOI or saline injection. (*Left*) Scatterplot comparing percentage of time mobile before and after DOI or saline administration. Average ± s.e.m values included. (*Right*) Average ± s.e.m change in percentage of total time mobile due to DOI or saline injections. Black dot marks example mouse used in A and Figure 1. (D) Same as B but for percentage of time spent in the center of the arena before and after injection of either DOI or saline. The center of the arena was defined as illustrated in A. (E) Same as B and C but for average pupil size across the recording before and after injection of either DOI or saline.

**Figure S2.**
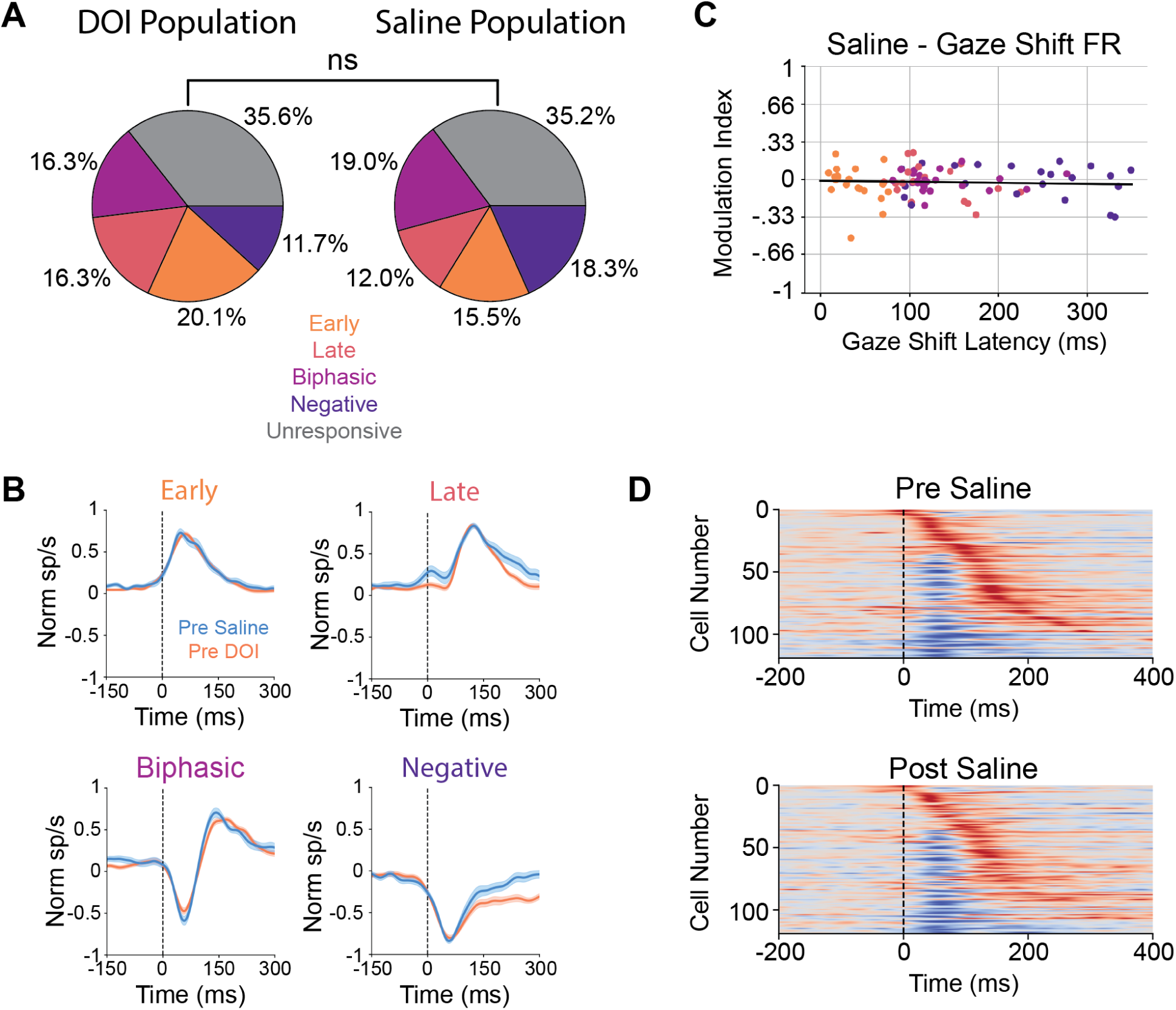
Saline does not significantly affect gaze shift responses. (A) Proportion of cells in each gaze shift cluster across DOI (*left*) and saline (*right*) datasets. (B) Average ± s.e.m. normalized pre-injection gaze shift response PETHs. (C) Gaze shift evoked firing rate modulation index vs gaze shift response latency for the saline population. Black line is a linear fit to the saline data. Compare to Figure 3D. (D) Temporal sequence of normalized gaze shift responses sorted along the y-axis by response latency before (*top)* and after (*bottom*) saline. In both plots cells are sorted by their pre-injection response latencies. Dotted line denotes gaze shift onset. Compare to Figure 3E. Latency values plotted in Fig 3F.

**Figure S3.**
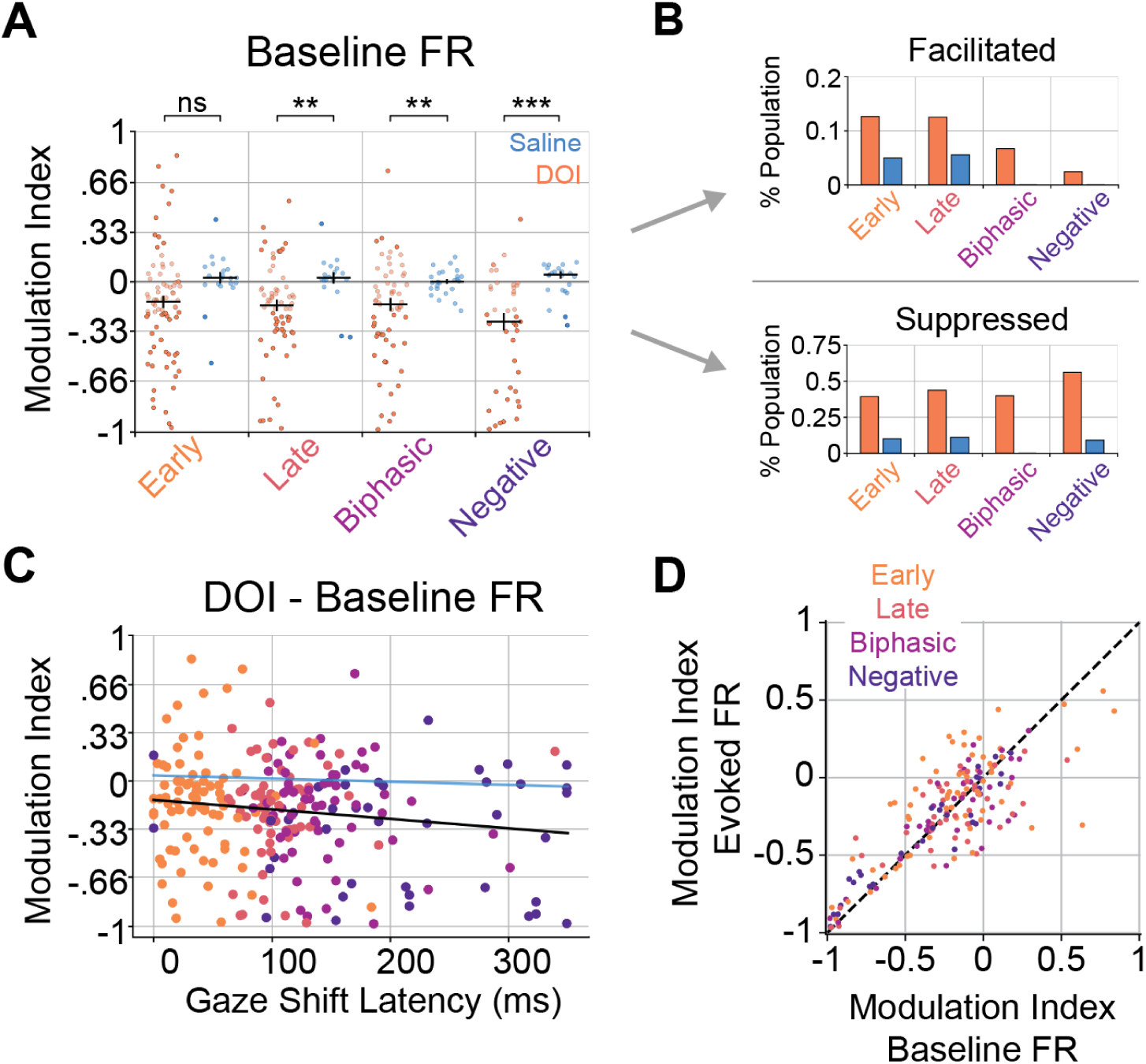
Cluster specific effects of DOI on baseline firing rates during free movement. (A) Modulation index of effect of DOI or saline administration on gaze shift baseline responses sorted by gaze shift cluster. Positive or negative values indicate injection-induced facilitation or suppression, respectively. Median ± s.e.m. values included for each condition shown in black. Values> 0.2 or < -0.2 were considered substantially modulated by the injection. Cells that were not substantially modulated were plotted as slightly transparent. (B) Percentage of each cluster that had baseline firing rate modulation indices > 0.2 (facilitated) or < -0.2 (suppressed) for DOI and saline datasets. (C) Baseline firing rate modulation index vs gaze shift response latency for the DOI population. Black line is linear fit to the DOI data, blue line is linear fit to the saline data (not plotted). (D) Scatterplot comparing the DOI-evoked modulation of baseline and gaze shift firing rates. The dotted line indicates y = x.

**Figure S4.**
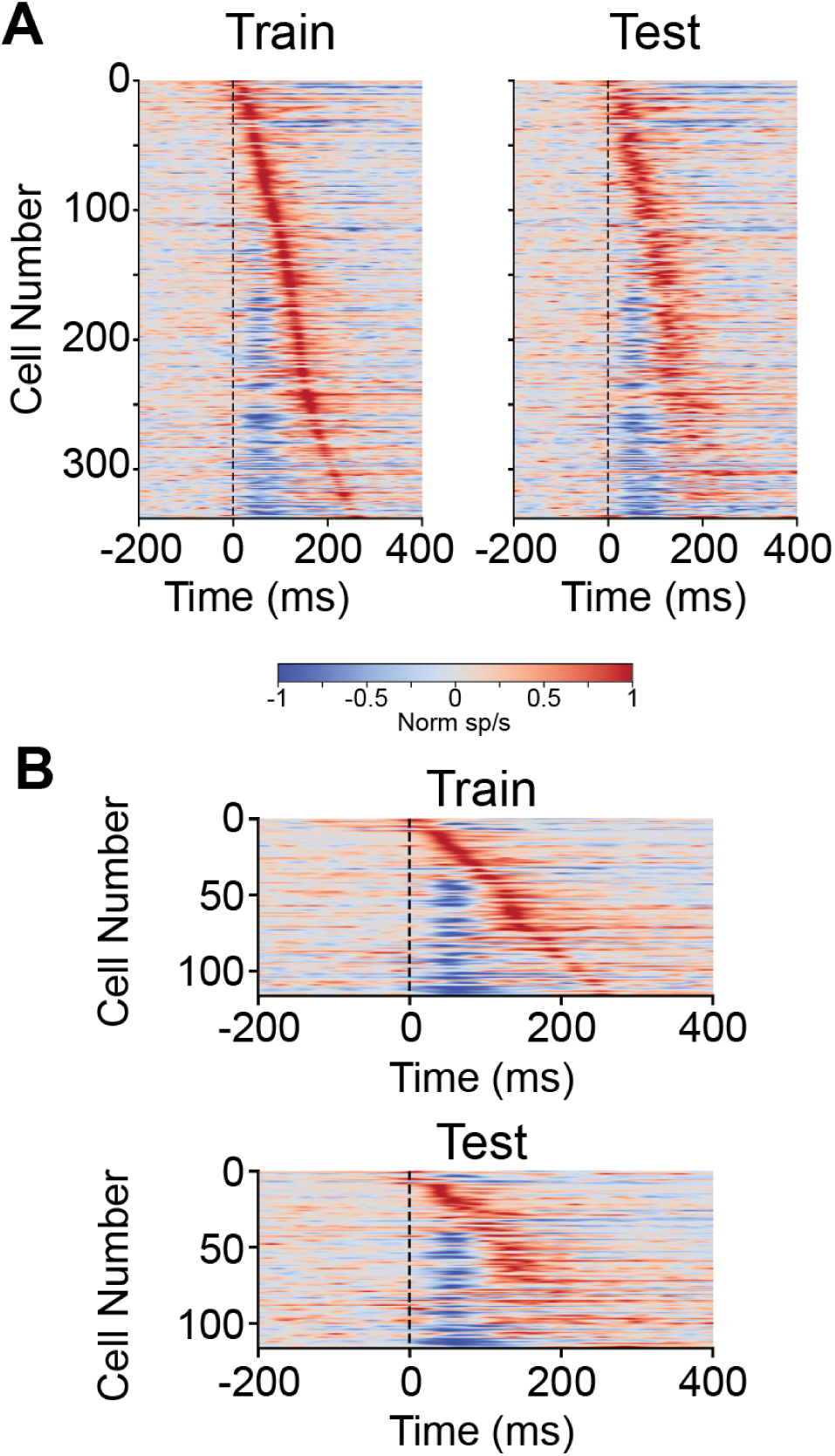
Cross-validation. (A) Cross-validation of pre DOI gaze shift PETHs for all responsive cells. Gaze shift times were randomly divided into two sets used to calculate PETHs in both the train (*left*) and test (*right*) sets. Both plots were sorted by the latency of the response in the train dataset. (B) Same as A but for the saline dataset.

**Figure S5.**
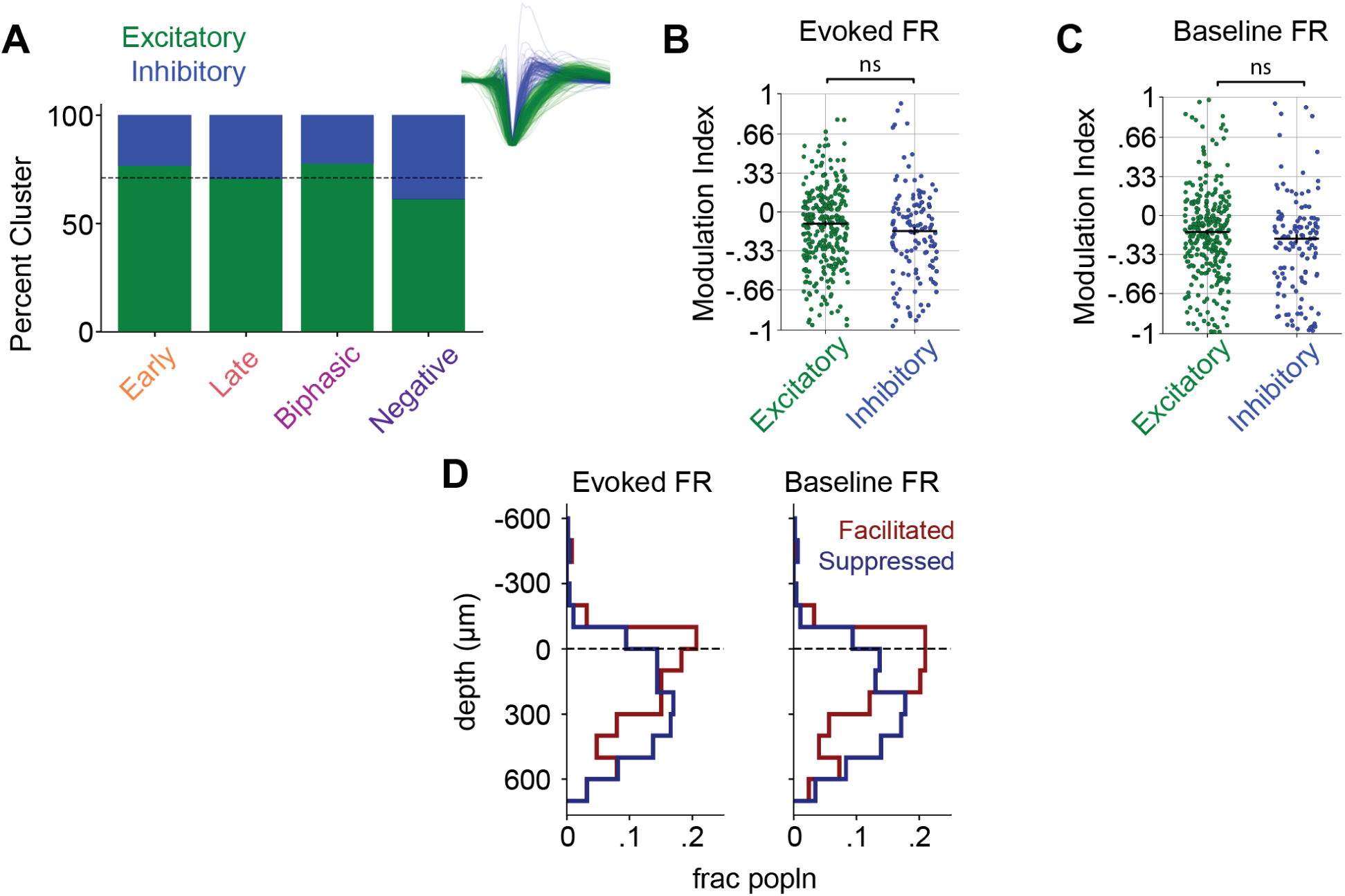
Effect of DOI on putative cell types and cortical lamina. (A) Fraction of putatively excitatory and inhibitory cells in each gaze shift response cluster. Excitatory and inhibitory groups were identified by k-means clustering on spike waveforms (shown to right). Dotted horizontal line denotes fraction of excitatory and inhibitory cells across the full population. (B) Gaze shift evoked firing rate modulation indices for DOI data sorted by waveform group. Median ± s.e.m. values included for each cluster (C) Same as B but for baseline firing rate. (D) Laminar depth of all cells sorted by whether DOI facilitated (red, modulation index > 0) or suppressed (blue, modulation index < 0) the gaze shift evoked (left) and baseline firing rate (right). Laminar depth was determined using the local field potential from multi-unit activity. Black dotted line (0 μm) is the estimated depth of cortical layer 5, to which depths were aligned.

## References

1. Shulgin, AT Shulgin, A (1991). Pihkal: a Chemical Love Story (Transform Press).

2. Mitchell, J.M., and Anderson, B.T. (2024). Psychedelic therapies reconsidered: compounds, clinical indications, and cautious optimism. Neuropsychopharmacology 49, 96–103. 10.1038/s41386-023-01656-7.

3. Lukasiewicz, K., Baker, J.J., Zuo, Y., and Lu, J. (2021). Serotonergic psychedelics in neural plasticity. Front. Mol. Neurosci. 14, 748359. 10.3389/fnmol.2021.748359.

4. Nardou, R., Sawyer, E., Song, Y.J., Wilkinson, M., Padovan-Hernandez, Y., de Deus, J.L., Wright, N., Lama, C., Faltin, S., Goff, L.A., et al. (2023). Psychedelics reopen the social reward learning critical period. Nature 618, 790–798. 10.1038/s41586-023-06204-3.

5. Carhart-Harris, R.L., Muthukumaraswamy, S., Roseman, L., Kaelen, M., Droog, W., Murphy, K., Tagliazucchi, E., Schenberg, E.E., Nest, T., Orban, C., et al. (2016). Neural correlates of the LSD experience revealed by multimodal neuroimaging. Proc. Natl. Acad. Sci. U. S. A. 113, 4853–4858. 10.1073/pnas.1518377113.

6. Hidalgo Jiménez, J., Kristjan Kaup, K., and Aru, J. (2026). Electrophysiological mechanisms of psychedelic drugs: A systematic review. Neurosci. Biobehav. Rev. 185, 106649. 10.1016/j.neubiorev.2026.106649.

7. Canal, C.E., and Morgan, D. (2012). Head-twitch response in rodents induced by the hallucinogen 2,5-dimethoxy-4-iodoamphetamine: a comprehensive history, a re-evaluation of mechanisms, and its utility as a model. Drug Test. Anal. 4, 556–576. 10.1002/dta.1333.

8. Halberstadt, A.L., van der Heijden, I., Ruderman, M.A., Risbrough, V.B., Gingrich, J.A., Geyer, M.A., and Powell, S.B. (2009). 5-HT(2A) and 5-HT(2C) receptors exert opposing effects on locomotor activity in mice. Neuropsychopharmacology 34, 1958–1967. 10.1038/npp.2009.29.

9. Arnt, J., and Hyttel, J. (1989). Facilitation of 8-OHDPAT-induced forepaw treading of rats by the 5-HT2 agonist DOI. Eur. J. Pharmacol. 161, 45–51. 10.1016/0014-2999(89)90178-7.

10. Horrocks, M., Mohn, J.L., and Jaramillo, S. (2025). The serotonergic psychedelic DOI impairs deviance detection in the auditory cortex. J. Neurophysiol. 133, 388–398. 10.1152/jn.00411.2024.

11. Michaiel, A.M., Parker, P.R.L., and Niell, C.M. (2019). A hallucinogenic serotonin-2A receptor agonist reduces visual response gain and alters temporal dynamics in mouse V1. Cell Rep. 26, 3475–3483.e4. 10.1016/j.celrep.2019.02.104.

12. Siegel, J.S., Subramanian, S., Perry, D., Kay, B.P., Gordon, E.M., Laumann, T.O., Reneau, T.R., Metcalf, N.V., Chacko, R.V., Gratton, C., et al. (2024). Psilocybin desynchronizes the human brain. Nature 632, 131–138. 10.1038/s41586-024-07624-5.

13. Wallace, D.J., Greenberg, D.S., Sawinski, J., Rulla, S., Notaro, G., and Kerr, J.N.D. (2013). Rats maintain an overhead binocular field at the expense of constant fusion. Nature 498, 65–69. 10.1038/nature12153.

14. Meyer, A.F., Poort, J., O’Keefe, J., Sahani, M., and Linden, J.F. (2018). A Head-Mounted Camera System Integrates Detailed Behavioral Monitoring with Multichannel Electrophysiology in Freely Moving Mice. Neuron 100, 46–60.e7. 10.1016/j.neuron.2018.09.020.

15. Michaiel, A.M., Abe, E.T., and Niell, C.M. (2020). Dynamics of gaze control during prey capture in freely moving mice. Elife 9. 10.7554/eLife.57458.

16. Parker, P.R.L., Martins, D.M., Leonard, E.S.P., Casey, N.M., Sharp, S.L., Abe, E.T.T., Smear, M.C., Yates, J.L., Mitchell, J.F., and Niell, C.M. (2023). A dynamic sequence of visual processing initiated by gaze shifts. Nat. Neurosci. 26, 2192–2202. 10.1038/s41593-023-01481-7.

17. Skyberg, R.J., and Niell, C.M. (2024). Natural visual behavior and active sensing in the mouse. Curr. Opin. Neurobiol. 86, 102882. 10.1016/j.conb.2024.102882.

18. Land, M. (2019). Eye movements in man and other animals. Vision Res. 162, 1–7. 10.1016/j.visres.2019.06.004.

19. Wallace, D.J., Voit, K.-M., Martin Machado, D., Bahadorian, M., Sawinski, J., Greenberg, D.S., Stahr, P., Holmgren, C.D., Bassetto, G., Rosselli, F.B., et al. (2025). Eye saccades align optic flow with retinal specializations during object pursuit in freely moving ferrets. Curr. Biol. 35, 761–775.e10. 10.1016/j.cub.2024.12.032.

20. Holmgren, C.D., Stahr, P., Wallace, D.J., Voit, K.-M., Matheson, E.J., Sawinski, J., Bassetto, G., and Kerr, J.N.D. (2021). Visual pursuit behavior in mice maintains the pursued prey on the retinal region with least optic flow. Elife 10, e70838. 10.7554/eLife.70838.

21. Meyer, A.F., O’Keefe, J., and Poort, J. (2020). Two Distinct Types of Eye-Head Coupling in Freely Moving Mice. Curr. Biol. 30, 2116–2130.e6. 10.1016/j.cub.2020.04.042.

22. Sharp, S.L., Shin, J., Martins, D.M., Jones, K., and Niell, C.M. (2025). Neural dynamics in superior colliculus of freely moving mice. bioRxiv. 10.1101/2025.04.16.648828.

23. Welch, A.C., Gonzales-Hess, N., Tarvin, T.A., Connor, J., and Smear, M.C. (2025). The impact of a psychedelic drug on olfactory search behavior by mice. bioRxiv. 10.1101/2025.07.09.663970.

24. Hegdé, J. (2008). Time course of visual perception: coarse-to-fine processing and beyond. Prog. Neurobiol. 84, 405–439. 10.1016/j.pneurobio.2007.09.001.

25. Skyberg, R., Tanabe, S., Chen, H., and Cang, J. (2022). Coarse-to-fine processing drives the efficient coding of natural scenes in mouse visual cortex. Cell Rep. 38, 110606. 10.1016/j.celrep.2022.110606.

26. Hénaff, O.J., Boundy-Singer, Z.M., Meding, K., Ziemba, C.M., and Goris, R.L.T. (2020). Representation of visual uncertainty through neural gain variability. Nat. Commun. 11, 2513. 10.1038/s41467-020-15533-0.

27. Marr, D. (2010). Vision: A Computational Investigation into the Human Representation and Processing of Visual Information (MIT Press).

28. Cassidy, C.M., Balsam, P.D., Weinstein, J.J., Rosengard, R.J., Slifstein, M., Daw, N.D., Abi-Dargham, A., and Horga, G. (2018). A perceptual inference mechanism for hallucinations linked to striatal dopamine. Curr. Biol. 28, 503–514.e4. 10.1016/j.cub.2017.12.059.

29. Grossberg, S. (2000). How hallucinations may arise from brain mechanisms of learning, attention, and volition. J. Int. Neuropsychol. Soc. 6, 583–592. 10.1017/s135561770065508x.

30. Carhart-Harris, R.L., and Friston, K.J. (2019). REBUS and the anarchic brain: Toward a unified model of the brain action of psychedelics. Pharmacol. Rev. 71, 316–344. 10.1124/pr.118.017160.

31. Klüver, H. (1966). Mescal, and mechanisms of hallucinations. -- (University of Chicago Press).

32. Bressloff, P.C., Cowan, J.D., Golubitsky, M., Thomas, P.J., and Wiener, M.C. (2002). What geometric visual hallucinations tell us about the visual cortex. Neural Comput. 14, 473–491. 10.1162/089976602317250861.

33. Schmack, K., Bosc, M., Ott, T., Sturgill, J.F., and Kepecs, A. (2021). Striatal dopamine mediates hallucination-like perception in mice. Science 372, eabf4740. 10.1126/science.abf4740.

34. Bauer, J., Margrie, T.W., and Clopath, C. (2025). Movie reconstruction from mouse visual cortex activity. 10.7554/elife.105081.1.

35. Jiang, Q., Shao, L.-X., Yao, S., Savalia, N.K., Gilbert, A.D., Davoudian, P.A., Nothnagel, J.D., Tian, G., Hung, T.S., Lai, H.M., et al. (2026). Psilocybin triggers an activity-dependent rewiring of large-scale cortical networks. Cell 189, 659–675.e22. 10.1016/j.cell.2025.11.009.

36. Shao, L.-X., Liao, C., Davoudian, P.A., Savalia, N.K., Jiang, Q., Wojtasiewicz, C., Tan, D., Nothnagel, J.D., Liu, R.-J., Woodburn, S.C., et al. (2025). Psilocybin’s lasting action requires pyramidal cell types and 5-HT2A receptors. Nature 642, 411–420. 10.1038/s41586-025-08813-6.

37. Weber, E.T., and Andrade, R. (2010). Htr2a gene and 5-HT(2A) receptor expression in the cerebral cortex studied using genetically modified mice. Front. Neurosci. 4. 10.3389/fnins.2010.00036.

38. Schmitz, G.P., Chiu, Y.-T., Foglesong, M.L., Magee, S.N., MacKinnon, M., König, G.M., Kostenis, E., Hsu, L.-M., Shih, Y.-Y.I., Roth, B.L., et al. (2025). Psychedelic compounds directly excite 5-HT2A layer V medial prefrontal cortex neurons through 5-HT2A Gq activation. Transl. Psychiatry 15, 381. 10.1038/s41398-025-03611-0.

39. Piscopo, D.M., El-Danaf, R.N., Huberman, A.D., and Niell, C.M. (2013). Diverse visual features encoded in mouse lateral geniculate nucleus. J. Neurosci. 33, 4642–4656. 10.1523/JNEUROSCI.5187-12.2013.

40. Gao, E., DeAngelis, G.C., and Burkhalter, A. (2010). Parallel input channels to mouse primary visual cortex. J. Neurosci. 30, 5912–5926. 10.1523/JNEUROSCI.6456-09.2010.

41. Al Shoyaib, A., Archie, S.R., and Karamyan, V.T. (2019). Intraperitoneal route of drug administration: Should it be used in experimental animal studies? Pharm. Res. 37, 12. 10.1007/s11095-019-2745-x.

42. Parker, P.R.L., Abe, E.T.T., Leonard, E.S.P., Martins, D.M., and Niell, C.M. (2022). Joint coding of visual input and eye/head position in V1 of freely moving mice. Neuron 110, 3897–3906.e5. 10.1016/j.neuron.2022.08.029.

43. Kleiner, M.B., Brainard, D.H., Pelli, D.G., Ingling, A., and Broussard, C. (2007). What’s new in Psychtoolbox-3. Perception 36, 1–16. 10.1068/v070821.

44. Niell, C.M., and Stryker, M.P. (2008). Highly selective receptive fields in mouse visual cortex. J. Neurosci. 28, 7520–7536. 10.1523/JNEUROSCI.0623-08.2008.

45. Senzai, Y., Fernandez-Ruiz, A., and Buzsáki, G. (2019). Layer-specific physiological features and interlaminar interactions in the primary visual cortex of the mouse. Neuron 101, 500–513.e5. 10.1016/j.neuron.2018.12.009.

46. Mathis, A., Mamidanna, P., Cury, K.M., Abe, T., Murthy, V.N., Mathis, M.W., and Bethge, M. (2018). DeepLabCut: markerless pose estimation of user-defined body parts with deep learning. Nat. Neurosci. 21, 1281–1289. 10.1038/s41593-018-0209-y.

47. Nath, T., Mathis, A., Chen, A.C., Patel, A., Bethge, M., and Mathis, M.W. (2019). Using DeepLabCut for 3D markerless pose estimation across species and behaviors. Nat. Protoc. 14, 2152–2176. 10.1038/s41596-019-0176-0.

